# An Accurate Bioinformatics Tool For Anti-Cancer Peptide Generation Through Deep Learning Omics

**DOI:** 10.1101/654277

**Authors:** Aman Chandra Kaushik, Mengyang Li, Dong-Qing Wei

## Abstract

The Anti-cancer targets play crucial role in signalling processes of cells. We have developed an Anti-Cancer Scanner (ACS) tool for identification of Anti-cancer targets in form of peptides. ACS tool also allows fast fingerprinting of the Anti-cancer targets of significance in the current bioinformatics research. There are tools currently available which predicts the above-mentioned features in single platform. In the present work, we have compared the features predicted by ACS with other on-line available methods and evaluated the performance of the ACS tool. ACS scanned the Anti-cancer target protein sequences provided by the user against the Anti-cancer target data-sets. It has been developed in PERL language and it is scalable having an extensible application in bioinformatics with robust coding architecture. It achieves a prediction accuracy of 95%, which is much higher than the existing tools.

## INTRODUCTION

Cancer is one of the most devastating diseases responsible for millions of mortalities worldwide. Different types of cancers are abundant in different countries, but lung cancer was reported as the most common cause of death in men while breast cancer was recorded in women. However, stomach, colorectal, liver and prostate cancers are more common in men. Whereas, cervix, lung, colorectal, stomach and breast cancer are uniformly distributed in women [1]. Different risk factors were reported to be associated with cancer but most common cause of this disease is the mutations in functionally significant somatic genes [2]. Due to high rate of morbidity and mortality, diagnosis, treatment and prevention of cancer is the utmost priority in the current area of research [3]. Global efforts revealed different approaches for the treatment of cancer including operational therapy; chemo agents-based treatment, hormonal, radiation and biological therapy. However, these approaches are constrained due to its high financial cost, adverse effects and low therapeutics output [4]. So, this conventional treatment has reduced the success rate of cancer treatment [5]. To tackle this devastating condition, new means of treatments are required. Recently, anti-cancer therapeutic vaccines have been developed and widely explored[6–10]. Anti-cancer vaccines (ACV), that contained short chain of amino acids usually less than 50 amino acids, were found to be exceptionally better than conventional chemotherapeutic agents [11]. Other advantages over conventional drugs includes specificity towards the target, no or less intracellular toxicity, alteration feasibility and high penetration power have made these vaccines as promising agents over the purposed methods. Due to promising output, such small vaccines have revolutionized the pharmaceutical markets and number of vaccine therapeutics have been increased in market [12]. Anti-cancer vaccines (ACV) and Antimicrobial vaccines (AMPs) are of same characteristics displaying similar properties. Like cellular surface negativity of bacteria, cancerous cells also possess negative charge and thus, both ACV and AMPs showed broad spectrum of activities. This negatively charge cells are of significant interest, making potential interaction with the cell surface and thus, selective toxicity can be achieved. These specificity properties divide these vaccines into two different categories; one that shows toxicity against all types of cells including bacterial, cancerous and normal cells while other depicts activity against only bacterial and cancerous cells [13–15].

Despite enormous therapeutic significance of Anti-cancer vaccines, till date, no tool for Anticancer vaccines and Anti-cancer proteins/peptides has been developed to identify the vaccines from wild and mutated cancerous protein sequences that can be used as anti-cancer vaccines. Large databases are available such as data from Clinical Proteomic Tumor Analysis Consortium (CPTAC)[16] which contains proteomic data from mass spectrometry analysis, and compare expression patterns of proteome and transcript. Also, Cancer Genome Atlas (TCGA)[17] is one of the reservoirs of cancerous datasets that holds RNA-sequence data sets. On the other hand, cancer transcriptomics data is provided by microarray expression analysis. Many different cancer mutations related data such as data from Cancer Genomics Hub[18], Catalogue of Somatic Mutations in Cancer (COSMIC) [19], SNP500Cancer [20], and UCSC Cancer Genomics Browser [21] are available.

Despite the availability of such huge data, no efforts have been made to analyze and retrieve important patterns from this big data that could be clinically significant for the treatment of cancer. Therefore, to support the scientific portfolio developing anti-cancer vaccines; we have developed Anti-Cancer Scanner (ACS) tool which accept the cancer associated proteins data including information from the above sources to analyze and retrieve important evidence from them and predict Anti-cancer vaccine from their target sequence using machine learning approach as shown in Figure 1. This proposed tool identifies vaccines/peptides from targeted cancerous protein which means it’s personalized vaccines for cancer patients. We believe that ACS will be helpful for both bioinformatics and experimental researchers working in the field of Anti-cancer vaccine based therapeutics.

**Figure 1.**
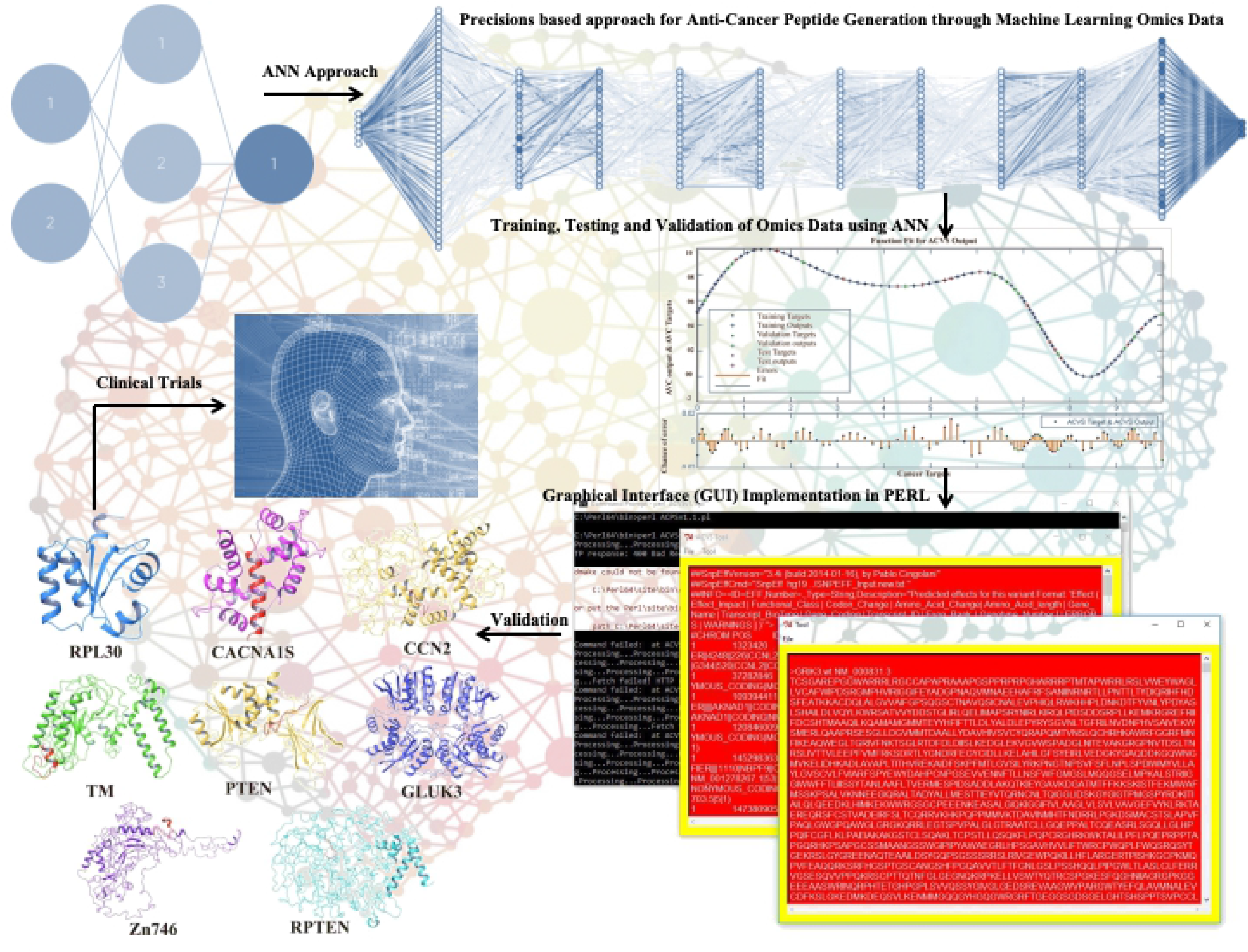
Schematic plan shows six steps employed of precisions based approach for anti-cancer peptide generation through machine learning omics data in form of agonist as promising strategy in the treatment of cancer

## METHODOLOGY

### Implementation

Following steps in the methodology were followed in the development of ACS:

#### Data mining of Anti-cancer targets

Many databases were used for data mining; many cancer databases are a molecular group of precise information scheme that gathers heterogeneous records of Anti-cancer targets.

#### Data set development of Anti-cancer targets

A dataset of Anti-cancer targets was developed, it includes 100 Anti-cancer targets, and input parameters used for ACS dataset were length of protein, origin character of mutation, mutated characters, and positions.

#### Development of algorithm for ACS

ACS is based on artificial neural network which is part of nature inspired algorithms that categorised into three important parts (Figure 2)

**Figure 2:**
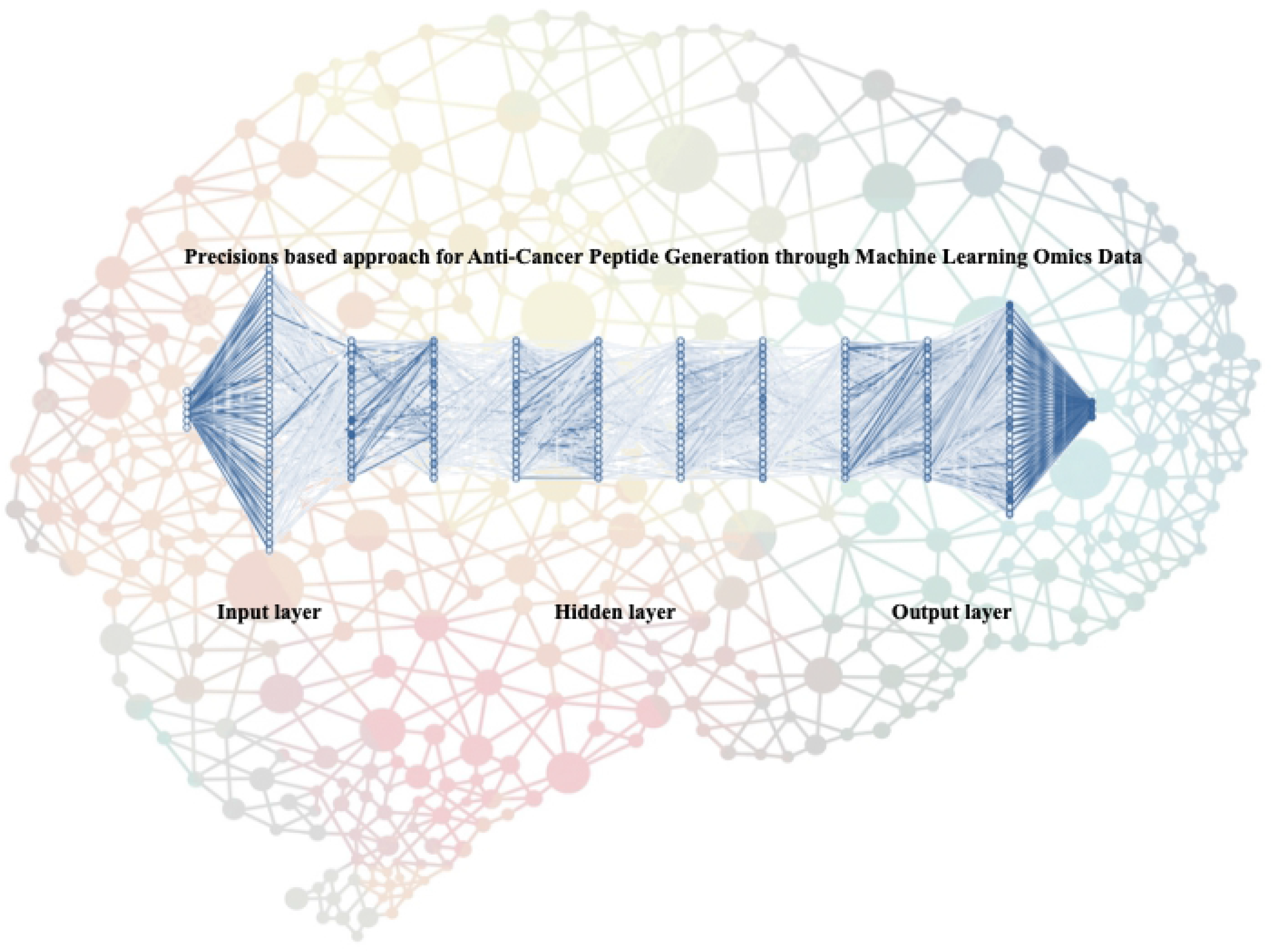
Neural network-based model consisting of connected submodels in where circles represent the input layer, hidden layer and output layer

1. Input layer of node
2. Hidden layer node and
3. Output node.

Each layer in neural network has one parallel node and consequently building the stacking. Output of each neuron is collective information of neurons multiplied by parallel weights with biased value while input value is transformed into output.

#### Normalization of Anti-cancer targets Dataset

Retrieves data for Anti-cancer targets dataset from various sources. Normalized dataset computed by V_new_ = (V_old_ – MinV) / (MaxV – MinV) * (D_max_ – D_min_) + D_min_.

Where, MinV is the minimum assessment of variable, MaxV maximum estimation of variable, D_max_ is the maximum estimation succeeding to normalization, D_min_ is the minimum assessment after normalization, V_new_ is new assessment after normalization and V_old_ assessment before normalization.

1. Input the data for training, and executed for training.
2. Set Anti-cancer targets network constraint.
3. Let learning count from 0;
4. Let learning count increase by 1;
5. Training stage iteration begins For ();
6. Input one sample then assign a new process and assign datasets.
7. Go to step 6 otherwise, (at least one dataset under the process).
8. Verify the sample to the stack for again test or training.
9. Iteration terminate when all samples have been tested
10. Learning error is computed.
11. Go to step 5, if the total learning error is 0.0
12. End
13. Calculate the neurons of output net_j_ = Σ_i=1~m_w_ji_ x_i_ +b_j_ where net_j_ is output neurons, w_ji_ connection weight neurons, x_i_ is input signal neurons and b_j_ is bias neurons.
14. Signal of output layers are calculated net_k_=TV_k_ + δ_K_^L^ where TV_k_ is target value of output neurons and δ_K_^L^ is the error of neuron.
15. Compute the error of neuron k. Step3 and Step6 are repetitive until SSE = Σ_i=1~n_(T_i_ – Y_i_)^2^, where T_i_ is actual assessment and Y_i_ is estimated assessment.

## RESULTS

ACS tool is very useful and reliable tool for Anti-cancer targets analysis. It generates maximum output using minimum input of Anti-cancer targets; using these applications users can determine the Anti-cancer vaccines. ACS tool which accept cancer associated proteins data including data from different cancerous resources; then analyze and retrieve important information from their cancer targets/proteins and predict Anti-cancer vaccine using machine learning approach. This proposed tool identifies vaccines/peptides from targeted cancerous protein which means it’s a personalized vaccine for cancer patients.

It is graphical user interface application (Figure 4) where user can easily import cancer target list with mutation information and ACS will predict optimized anti-cancerous peptides from those imported targets using artificial neural network; get retrieves the data for anti-cancer targets dataset and fiddle with defined series of attributes and avoid the infiltration of neurons. Input the cancer target data (cancer proteins) list for training, interrelated values of input cancer data and output are executed for training where anti-cancer targets network constraint like length of protein, origin character of mutation, mutated characters, and positions which is numeral of hidden layers (4 hidden layers produce better union). Assigning a new process and put new knowledge datasets under this process and analyze the final output of artificial neural network, and verify the sample to stack again for test or training. Calculate the neurons of output, every neuron output signals are calculated shown in Figure 3.

**Figure 3:**
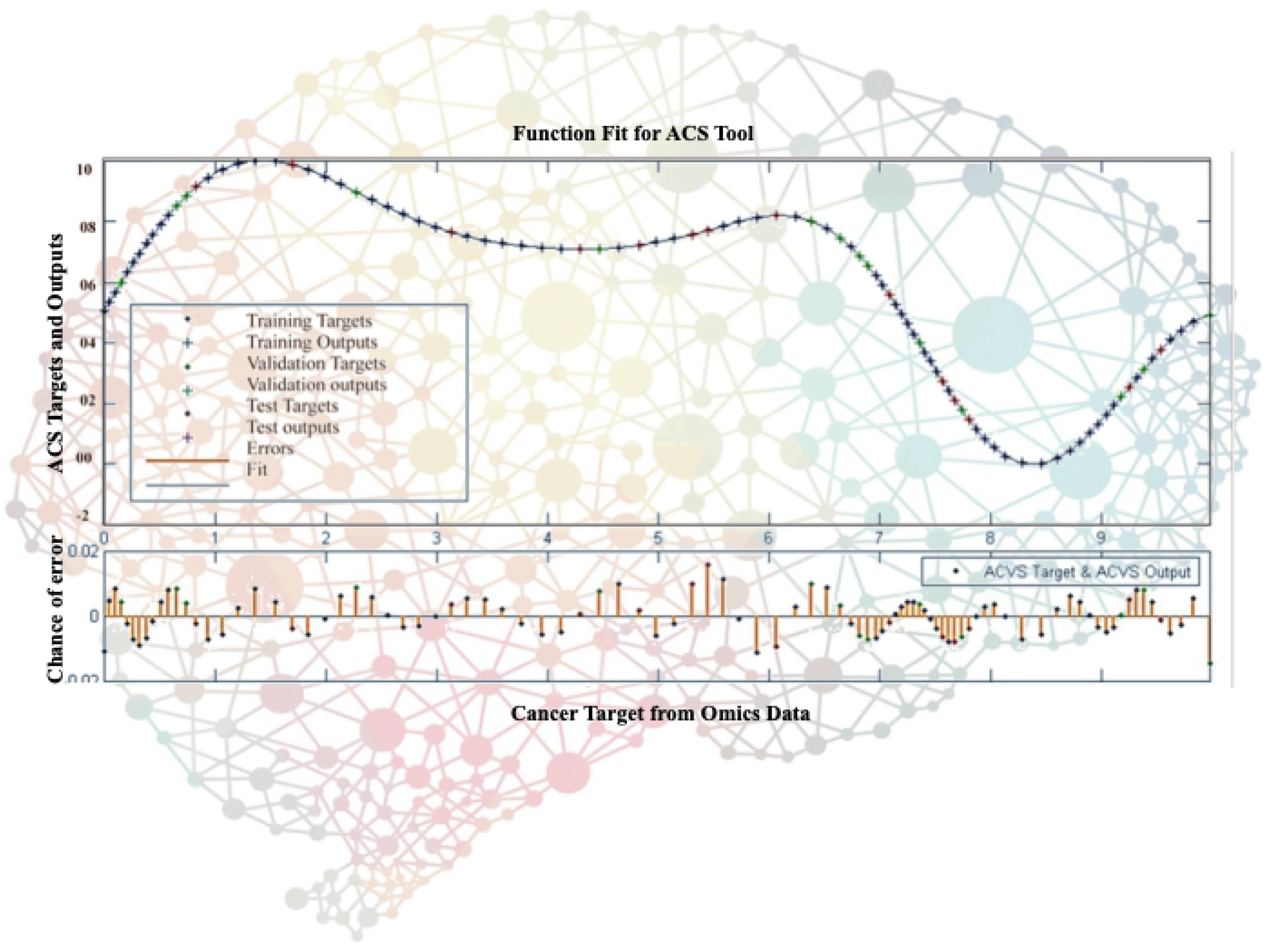
ACS performance. Graph represents the training, output and validation of targets; test output of targets; chance of error and function fit using artificial neural network approach, where upper X axis represents function fit and Y axis represents the ACS output and target dataset performance while lower X axis represents input data and Y axis represents the chance of performance

### Overview of the procedure

#### Input/output Instruction

1. Enter the sequence accession number. Many formats can be used: Raw/Plain, EMBL, Pearson/Fasta, etc. for further analysis.
2. Submit the job by clicking on submit button.
3. After you submit your query, a set of scores will be calculated, depending on which option you selected. Based on the scores, the vaccine will be ranked.

#### Peptide Motif Search

This package allowed users to locate and rank 8-mer, 9-mer, or 10-mer peptides that contain peptide binding motifs.

#### Input Instructions

1. Choose the gene sequence of interest from list with accession number through menu button. The selection of ID determines which coefficient table in scoring program will be used on the selected sequence.
2. Choose the length of subsequences, program then extracted the information for submitted input sequence and calculated the scores and ranks.
3. Choose whether to display the submitted input sequence on the output page using button.
4. Submit the job by using submit button.

#### Output Page Returned to the User

Following query submission, a set of scores are calculated for all 8-mer, 9-mer, or 10-mer subsequences contained in the input sequence, depending on selected option. Based on the scores, subsequences are ranked. This task is supposed to complete within a few seconds. After, a display page then returned with following information;

1. A short table of user-input parameters (for verification and later recall), along with some scoring data and other useful info (includes number of scores calculated, number of scores requested, number of scores reported back in the output table, and length of users input sequence)
2. The score output table, where the results of calculations on subsequences are displayed.
3. A listing of submitted input peptide sequence (if you have asked for the sequence to be echoed back)
4. Each row of the score output table consists of four columns. The values for these items represents the ranking of subsequence. The starting position in query protein sequence for the first amino acid residue of subsequence. A residue list 8-mer, 9-mer, or 10-mer peptide subsequence itself.

An estimated numerical score for subsequence (upon which the rank in the first column is based) shown in Figure 4.

**Figure 4:**
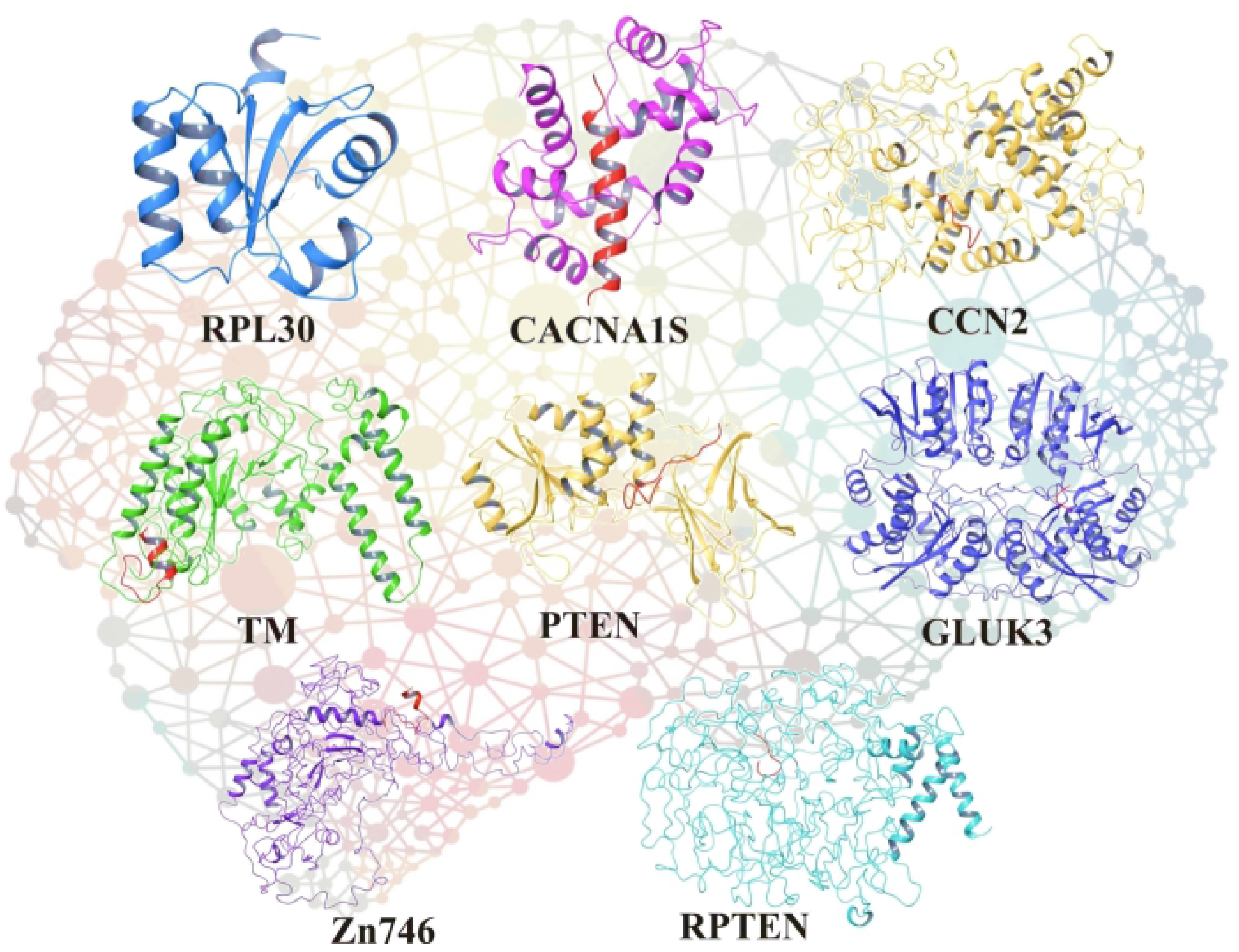
Front view of Anti-Cancer Scanner Tool through Machine Learning Omics Data **Validation**. For the validation of ACS tool, we took ten random cancer targets from dataset as a target and ten ACS generated optimized cancer vaccines/peptides (outputs) against same cancer targets, and applied molecular docking for each cancer targets and ACS suggested peptides (**Figure 5 and Table 1**).

**Table 1.**
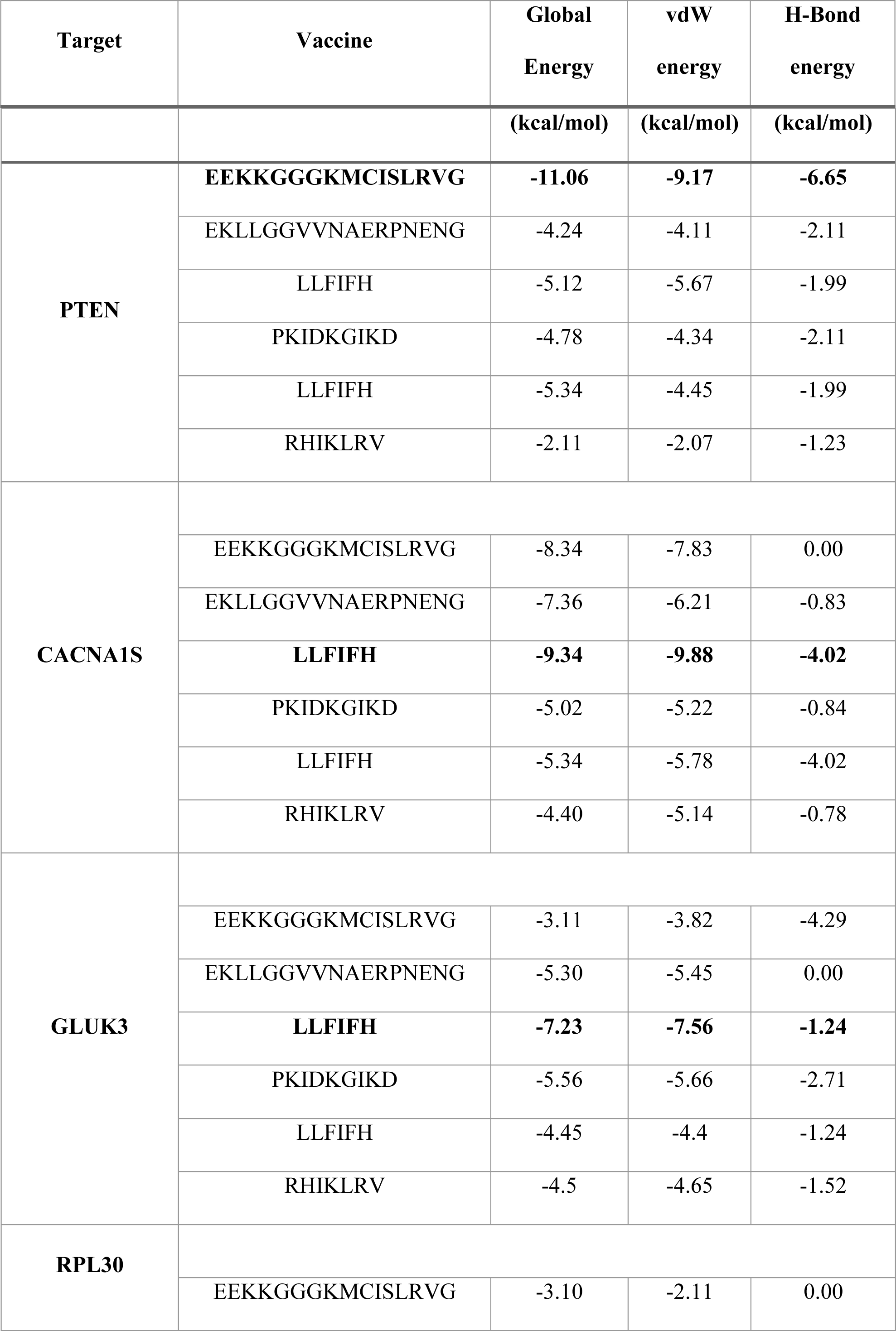
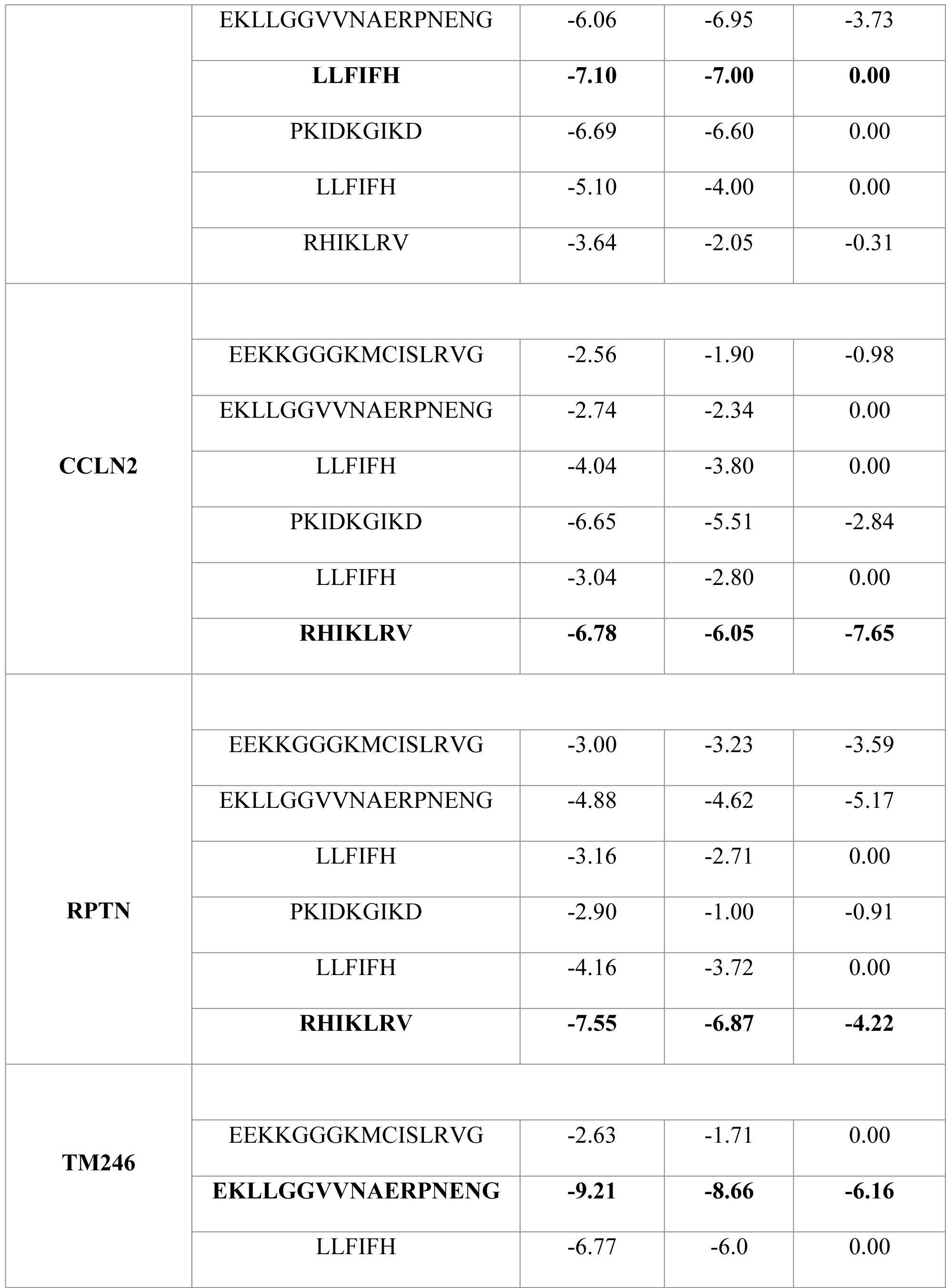
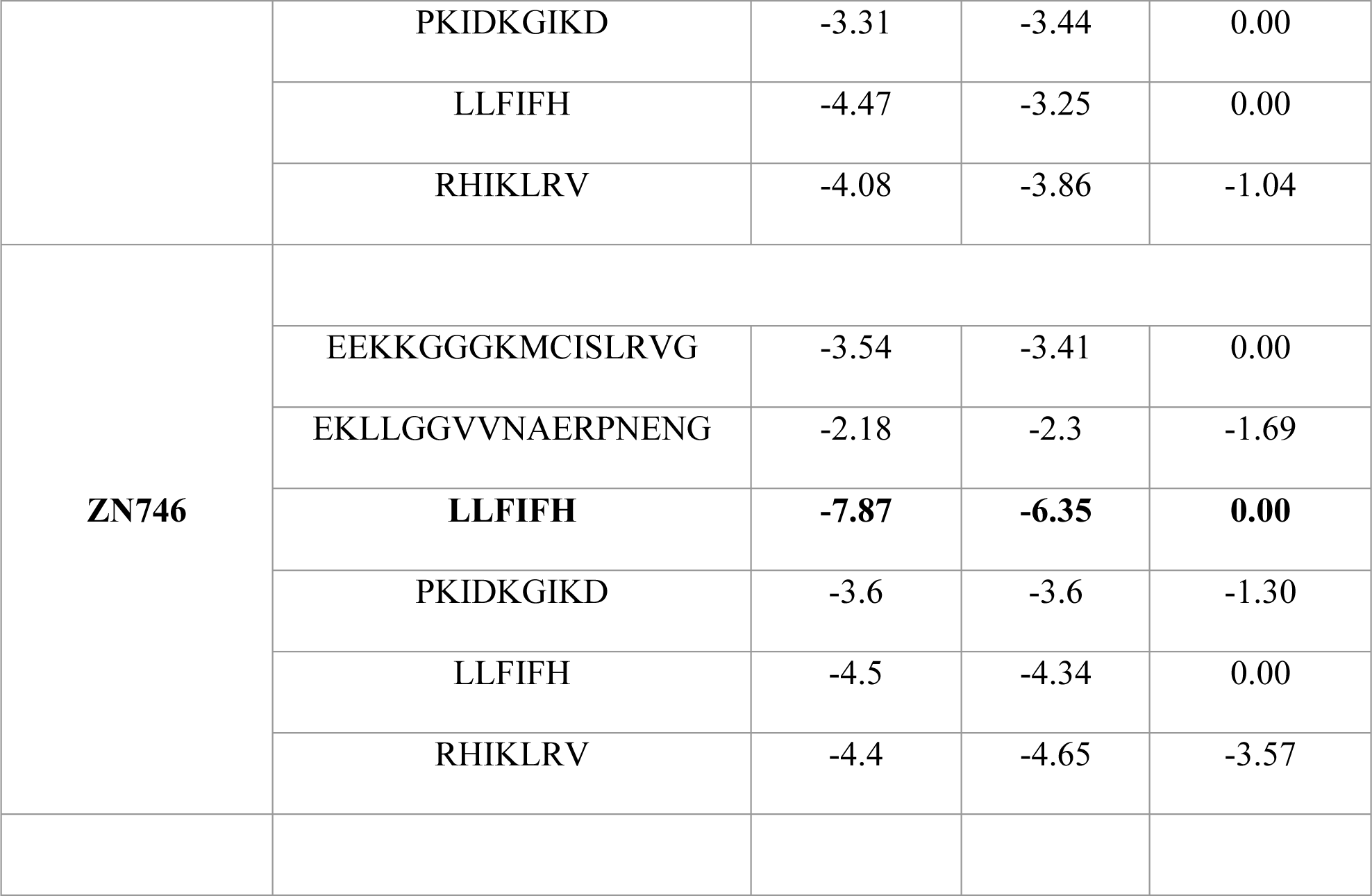
AutoDock energy for the best ranked complexes where top table represents the cancer targets, ACS suggested vaccines/peptides, global energy in kcal/mol, vdW energy in kcal/mol and H-Bond energy in kcal/mol of cancer targets and ACS suggested vaccines/peptides.

#### ACS tool advantages and applications

ACS is based on machine learning approch for identification of Anti-cancer targets based on their binding affinity. There is no such type of tool for identifying or classifying Anti-cancer targets. The advantage of ACS is that it analyses information for Anti-cancer targets which can be further assisted in precision drug discovery, identification of novel drug targets, drug designing for these novel targets as shown in Figure 6.

**Figure 5:**
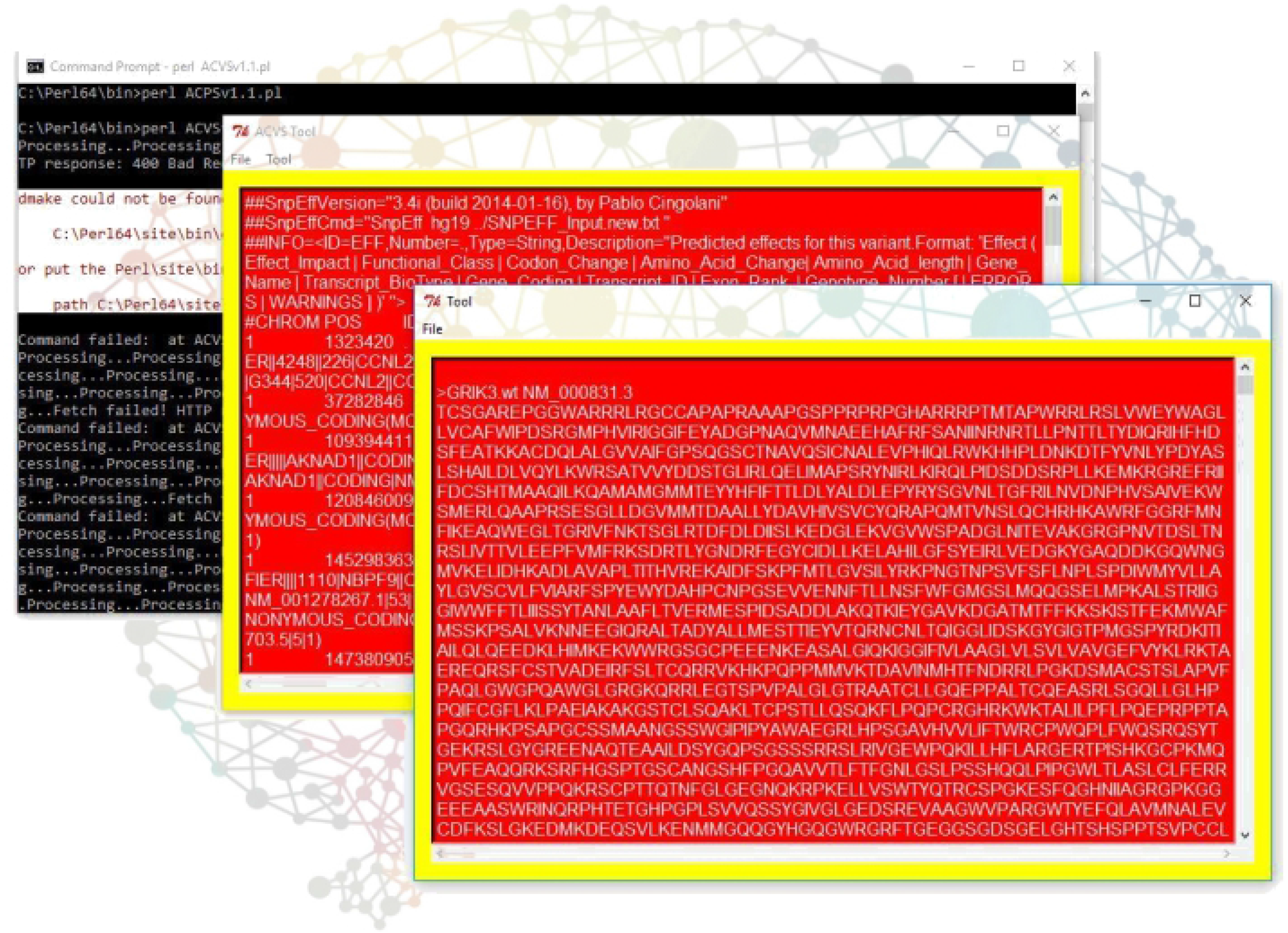
The figure is showing the interaction predicted peptides from their respective protein sequences. AutoDock were used to dock and define the interactions

**Figure 6.**
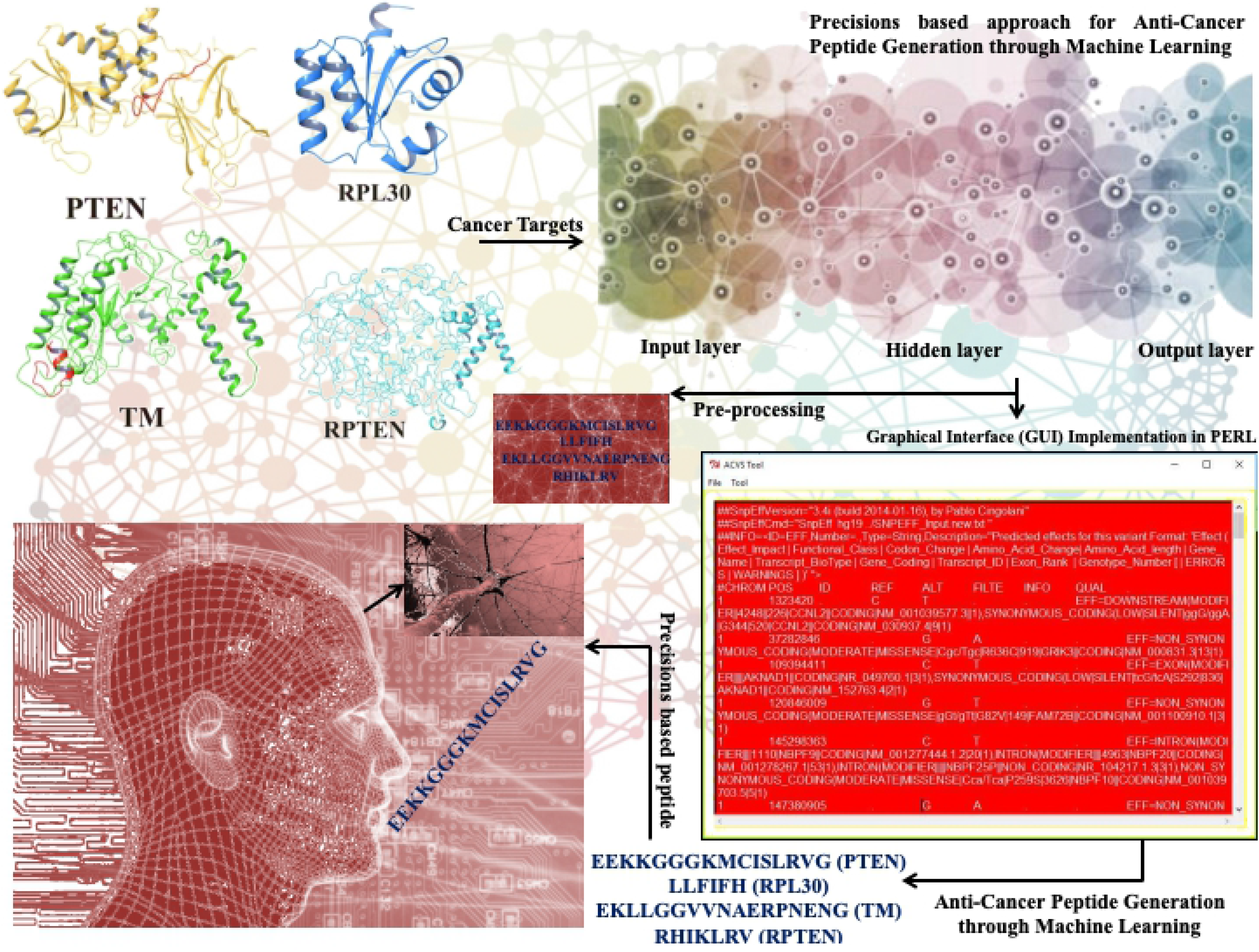
Schematic plan depicts the employed of precisions based peptides for anti-cancer peptide generation through machine learning

## CONCLUSION

We have proposed machine learning algorithm to identify Anti-cancer targets based on their binding affinity and implemented in artificial neural network. There is no such type of tool for identifying or classifying Anti-cancer targets using artificial neural network. It is graphical user interface application where user can easily import cancer target list with mutation information and ACS will predict optimized anti-cancerous vaccines/peptides from those imported targets using artificial neural network; get retrieves the data for anti-cancer targets dataset and fiddle with defined series of attributes and avoid the infiltration of neurons. ACS tool is advantageous for finding information about Anti-cancer targets which can be further assisted or employed in precision drug discovery, identification of novel drug targets, drug designing for these novel targets with a prediction accuracy of 95%. Black box approach were used for ACS script testing, where scripts are examining the functionality of ACS tool. ACS also inbuilt white box testing, where users opposed the functionality of ACS.

## LIST OF ABBREVIATIONS

ACV: Anti-cancer vaccines
AMPs: Antimicrobial vaccines
CPTAC: Clinical Proteomic Tumor Analysis Consortium
TCGA: Cancer Genome Atlas
COSMIC: Catalogue of Somatic Mutations in Cancer
ANN: Artificial neural network
EMBL: European Molecular Biology Lab

## ACKNOWLEDGMENTS

The simulations in this work were supported by the Center for High Performance Computing, Shanghai Jiao Tong University.

## AUTHORS CONTRIBUTIONS

Conceptualization: DQW, ACK

Data curation: ACK

Formal analysis: ACK

Funding acquisition: DQW

Investigation: DQW, ACK

Methodology: ACK

Project Administration: DQW

Resources: ACK, DQW

Supervision: DQW

Validation: DQW, ACK, ML

Writing-original draft: ACK

Writing-review draft: ACK, ML

## AVAILABILITY OF DATA AND MATERIAL

The data during and/or analyzed during the current study available from the corresponding author on request.

## ETHICS APPROVAL AND CONSENT TO PARTICIPATE

Not applicable.

